# The human olfactory bulb process odor valence representation and initiate motor avoidance behavior

**DOI:** 10.1101/2021.01.20.427468

**Authors:** Behzad Iravani, Martin Schaefer, Donald A. Wilson, Artin Arshamian, Johan N. Lundström

**Affiliations:** Department of Clinical Neuroscience, Karolinska Institutet, 17177 Stockholm, Sweden; Nathan Kline Institute for Psychiatric Research, Orangeburg, NY 10962, USA; Department of Child and Adolescent Psychiatry, New York University Langone Medical School, New York, NY 10016, USA; Monell Chemical Senses Center, Philadelphia, PA 19104, USA; Department of Psychology, University of Pennsylvania, Philadelphia, PA 19104, USA; Department of Psychology, Stockholm University, 10405 Stockholm, Sweden; Stockholm University Brain Imaging Centre, Stockholm University, 11415 Stockholm, Sweden

## Abstract

Determining the valence of an odor to provide information to guide rapid approach-avoidance behavior is thought to be one of the core tasks of the olfactory system, yet little is known of its initial neural mechanisms or subsequent behavioral manifestation in humans. In two experiments, we measured the functional processing of odor valence perception in the human olfactory bulb (OB)—the first processing stage of the olfactory system—using a non-invasive method as well as assessed subsequent motor avoidance response. We demonstrate that odor valence perception is associated with both gamma and beta activity in the human OB. Moreover, we show that negative, but not positive, odors initiate an early beta response in the OB, a response that is linked to a preparatory neural motor response in motor cortex. Finally, in a separate experiment we show that negative odors trigger a full-body motor avoidance response, manifested as a rapid leaning away from the odor, in the time period predicted by the OB results. Taken together, these results demonstrate that the human OB processes odor valence in a sequential manner in both the gamma and beta frequency bands and suggest that early processing of unpleasant odors in the OB might underlie rapid approach-avoidance decisions.

## INTRODUCTION

Survival of any organism is dependent on approach-avoidance mechanisms; avoiding dangerous- and approaching rewarding stimuli. Among our senses, the olfactory system seems specifically tuned to aid approach-avoidance decisions and in particular, to assist in avoiding potentially dangerous stimuli. It is not surprising then that the very first stage of the central olfactory system, the olfactory bulb (OB), processes various information directly related to whether an odor should be avoided (Kay and Laurent 1999).

In non-human animals, the OB demonstrates rapid plasticity to aversive stimuli (Kay and Laurent 1999) and has dedicated processing of odors innately associated with threats (Kobayakawa et al. 2007). Sensory systems are normally attuned to signals indicating negative outcomes for the individual given that a failure to respond to such stimuli may lead to fatal consequences (Haselton and Nettle 2006). For example, fast responses are arguably more important in respect to initiating the action to withdraw from toxic fumes than the need for speed to initiate the action to approach positive odor sources. The perceptual equivalent to the motor-driven approach-avoidance system in the olfactory system is the subjective perceptual experience of an odorant’s valence. Here, perceived unpleasantness of odorants emitted from potentially dangerous sources, such as for example rotten food, is translated to avoidance (Yeshurun and Sobel 2010). However, the underlying neural mechanism for this system is largely unknown. There are two major reasons for this. First, it is difficult to assess the subjective experience of a novel odorant’s valence in animal models. Second, although assessing subjective measurements from humans is straight-forward, until recently, there has been no method that allows a non-invasive measure of neural signals from the human OB. With that said, several brain imaging studies on humans have targeted the central processing mechanisms underlying valence perception. Here, valence perception has mainly been localized to the orbitofrontal cortex, OFC (*cf*. Seubert et al. 2017). However, the OFC is an area that is situated relatively late in the central olfactory processing stream (Mainland et al. 2014) and neural processing of odor avoidance in non-human animals have been localized much earlier in the processing stream, already one synapse away from the odor receptors, in the OB (Doucette et al. 2011; Kermen et al. 2016). Thus, it is necessary to study the OB to establish the underlying neural mechanism of the earliest processing stages to understand how the olfactory system process the subjective valence of an odorant, the first stage of an approach-avoidance decision.

Based on past studies in non-human animals, we hypothesized that the OB in awake humans would demonstrate early valence-differential processing and induce a preparatory motor approach/avoidance response according to perceived odor valence. In Experiment 1, we determined whether odor valence is processed by the human OB by means of a recently developed method that allows a direct, but non-invasive measurement of the human OB (Iravani et al. 2020). We found that subjective odor valence could be linked to gamma and beta activity in the human OB, independent of respiration, and that an early beta activity in OB processing was linked to motor cortex processing in a valence-dependent manner. Based on these results, in Experiment 2, we assessed whether humans, akin to non-human animals (Arshamian et al. 2017), demonstrate a rapid, full-body approach/avoidance response to odors in a valence-dependent manner in the time-period predicted by Experiment 1. We found that participants rapidly moved away from a negative odor source. Interestingly, only unpleasant odors produced a consistent motor response and, importantly, this response aligned temporally with the valence-associated activity in the OB demonstrated in Experiment 1.

## RESULTS

### Early phase amplitude coupling between beta and gamma in OB

Odor-evoked neural signals in response to 6 odors with varying valence were recorded from 4 electrodes located directly above the eyebrows, which, in combination with 64 EEG scalp electrodes, were used to extract source space electrobulbogram (EBG) (Iravani et al. 2020) signals from the OB (**Figure 1a**). Inhalation phase-locked odor stimuli were delivered using a sniff triggered, computer controlled, and temporally precise olfactometer (Lundström et al. 2010). Odor delivery delay (∼200ms) was measured with a photoionization detector and adjusted for in all analyses (Ohla and Lundström 2013). After each odor stimulus, participants rated perceived odor intensity, valence, and familiarity. A total of 19 participants participated in 3 separate and seemingly identical sessions, comprising a total of 540 trials per participant. Next, we removed trials with artifacts including muscle and blink (see the method section for details) by which on the average, 27.92 ± 10.49 clean trials per odor were included in analysis for each individual. Hence, considering all 6 odors, the total number of trials for each individual included in our analyses was on average 167.52 ± 25.81. More importantly, there was no statistical difference between the number of trials across odors, *F*(5,108) = .39, *p* > .86, indicating that after the artifact rejection, the experimental design remained balance.

**Figure 1.**
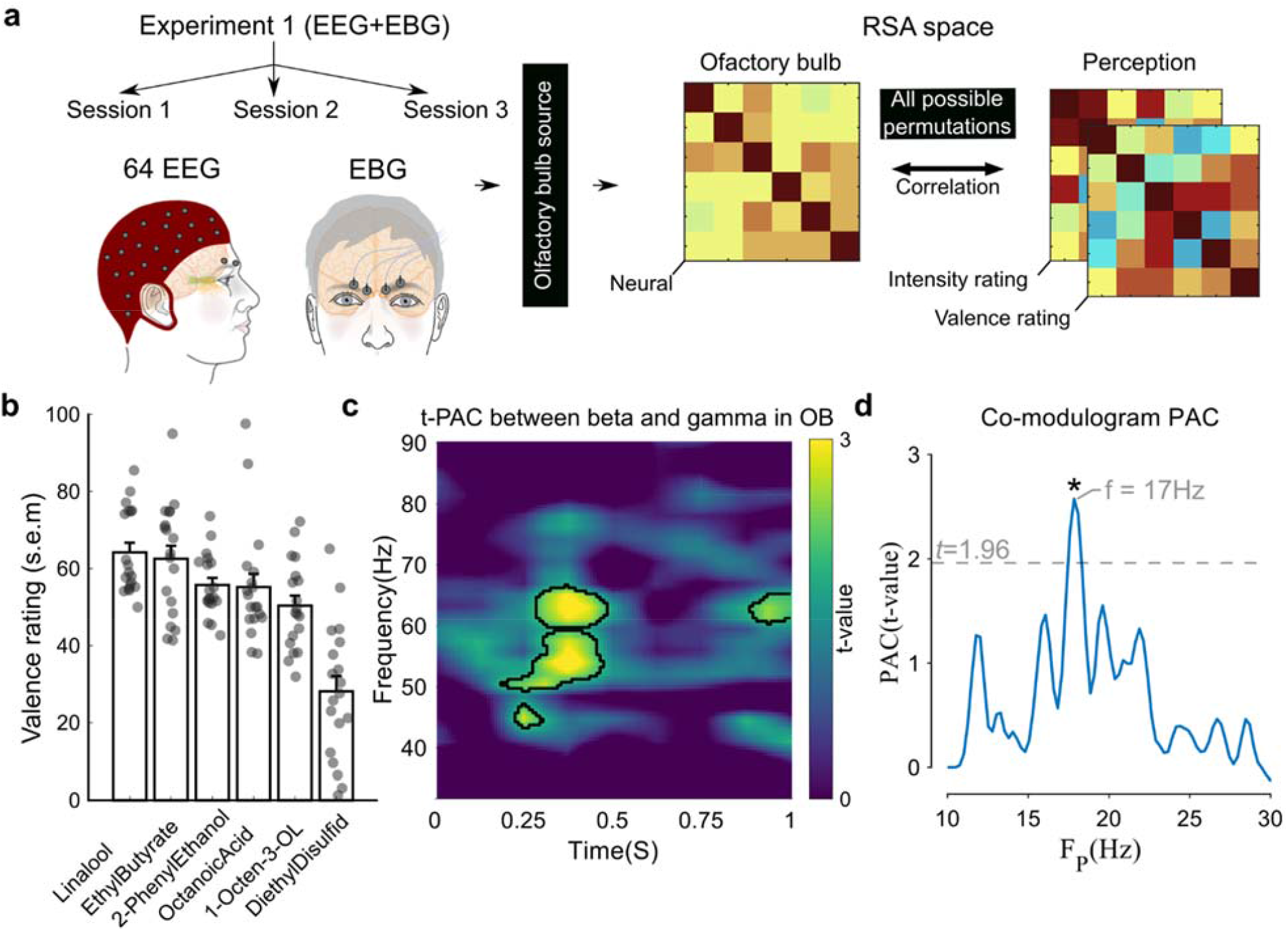
Early phase amplitude coupling between beta and gamma in the OB. **a**) Methodological summary of Experiment 1 where individuals (n=19) were tested during three separate sessions that were subsequently merged. Source reconstruction was performed using the EEG/EGB electrodes in combination with multi-spherical head model and digitalized electrode positions to extract olfactory bulb time-course. Representation dissimilarity matrixes were constructed for both olfactory bulb neural signals and perceptual ratings; subsequent partial Pearson correlations were derived for each time point from all possible permutations. **b**) Group mean perceived valence ratings of the 6 odors in Experiment 1. Individual’s mean ratings are indicated with filled circles and show a large variability between participants in rated valence of each odor. Note that for analyses, valence ratings of each individual were used to create a common structure with the DISTATIS method (Abdi et al. 2005). Error bars represent standard error of the mean (s.e.m). **c**) Heat map showing the strength of phase amplitude coupling as function of time. Compared to background, a significant coupling around 53-65 Hz (significant results assessed with permutation testing and marked with black boundaries) starts around 250 ms after odor onset. **d**) The co-modulogram between the beta and gamma bands (∼53-65 Hz) during the whole 1s indicates that the coupling appeared in beta band around ∼16-18 Hz. Significant peak marked with asterisk and assessed with student t-test. The statistical threshold for detecting significance (t = 1.96 equal to p < 0.05) is marked with gray dashed line. FP denotes frequency of slower oscillation or frequency phase.

We have previously established that the EBG measure is a valid and reliable measure (Iravani et al. 2020) but prior to our main analysis, we estimated the quality and spatial dispersion of the reconstructed OB signal within this unique dataset. To this end, we used a simulation where the spatial dispersion of three levels of signal-to-noise-ratio were assessed, namely a hypothetical ideal, the empirical level, and two-fold lower (i.e. two-fold larger noise level) than empirical level. This analyses confirmed that our source reconstruction method can successfully isolate OB’s EBG signal in source space given that the spatial gain was similar to the hypothetical ideal condition when assessing a signal-to-noise-ratio similar to what we empirically observed in the current dataset (**Figure S1**).

To allow direct comparisons between neural and behavioral data, we used Representational Similarity Analysis (RSA)—a multivariate method that compares similarity (e.g. correlation) matrixes between continuous relationships to determine the representational geometry on the individual level (Kriegeskorte et al. 2008), therefore allowing direct comparisons between different parameters without being hindered by difference in scaling and other inherent differences between measuring techniques. In this case, we assess how well a perceptual feature can be decoded from neural activity (presented as degree of similarity between measures’ Representation Dissimilarity Matrix, RDM). The perceptual and neural population RDMs were initially derived on the individual level as relationship-distances between individual odors, separately for perceptual and neural space, and later assessed for similarities between them in group level analysis (Abdi et al. 2005). In other words, for each time-point in the OB recording, we assess whether the relationship between the 6 tested odors in the neural space are similar to the relationship between ratings of same odors in the perceptual space. Statistical relationships are then assessed for the group. Specifically, we compared how odor-induced neural activity in the OB within the gamma and beta bands corresponded to individual valence ratings of the same odors (**Figure 1b**).

We initially assessed the relationship between activity of the beta and gamma frequency bands, determining the frequencies coupled together in OB and later the activity of these frequencies to perceived valence in RSA. Phase-amplitude coupling (PAC), a subclass of cross frequency-coupling phenomena, has been identified as a neural mechanism detectable in most mammals and critical for information processing in a multitude of brain regions (Buzsáki et al. 2003; Bragin et al. 1995; Kendrick et al. 2009; Lakatos et al. 2005; Axmacher et al. 2010; Cohen et al. 2009). Here, the phase of the lower frequency oscillation drives the power of the coupled higher frequency oscillation. Different functional roles have been attributed to PAC, including sensory signal detection (Händel and Haarmeier 2009), executive functions (Tort et al. 2009), and attentional selection (Schroeder and Lakatos 2009). Specifically for olfaction, it has recently been demonstrated that PAC in the OB shapes early sensory processing in mice (Losacco et al. 2020). Given this, we examined PAC between beta and gamma oscillation within the OB and its relation to the processing of the individual’s odor valence using RSA. To gain a temporal dimension of the PAC, we used time-resolved phase-amplitude coupling (t-PACSamiee et al. (2017)), a method that also incorporates the temporal dynamic of the signal.

As a first step, we assessed the relationship between beta and gamma bands with t-PAC within the first second after receiving an odor stimulus to determine whether the frequency of bands demonstrate phase-amplitude coupling and, if so, at which frequencies. We found significant PAC between beta and gamma already at 250ms after odor onset (∼53-65 Hz), as assessed by Monte Carlo permutation test (**Figure 1c**; *t* = 3.85, *p* < .006 and *CI* = [.002, .006]). Next, we assessed the comodulogram between beta and the detected range in gamma oscillations to isolate frequencies of interest in the beta band. We found that this coupling operates in the beta band within 16 to 18Hz (*t* = 2.57, *p* < .009, *CI* = [0.002, 0.012]; **Figure 1d**). The t-PAC and the co-modulogram results guided our subsequent neuronal and valence RSA analysis by isolating signal of interest.

### Early gamma and late beta activity relate to perceived odor valence

Band-passed OB reconstructed time courses were transformed into a complex signal using Hilbert transform. Both amplitude, as well as phase were used to construct neural and perceptual valence RDMs. This was performed at each time point separately for each frequency band (gamma/beta) using the Euclidean distance across 6 odors that varied in valence, resulting in a sequence of RDMs (**Figure 1a**). Then, maximum partial Pearson correlations were calculated by an approximate 80ms wide non-overlapping sliding window, sweeping 0 to 1s after odor onset anchored to inhalation, between two sequences of RDMs (**Figure 2a**). This resulted in a correlation time course while controlling for perceived intensity (**Figure 2b**). To test the significance of the correlation at each time point, all possible permutations (n=720) were tested and exact *p*-values were computed. We found time-points of significant associations in RSA space between sub-band of gamma activity (53-65 Hz, consequent to PAC result) and perceived odor valence around 250-325ms (*r*_*1*_ = .60, *p*_*1*_ < .010, *CI* = [.56, 1]) (**Figure 2b**; adjusted for measured olfactometer delay). The distribution and exact *p*-value for the significant instance is shown in **Figure 2c**.

**Figure 2.**
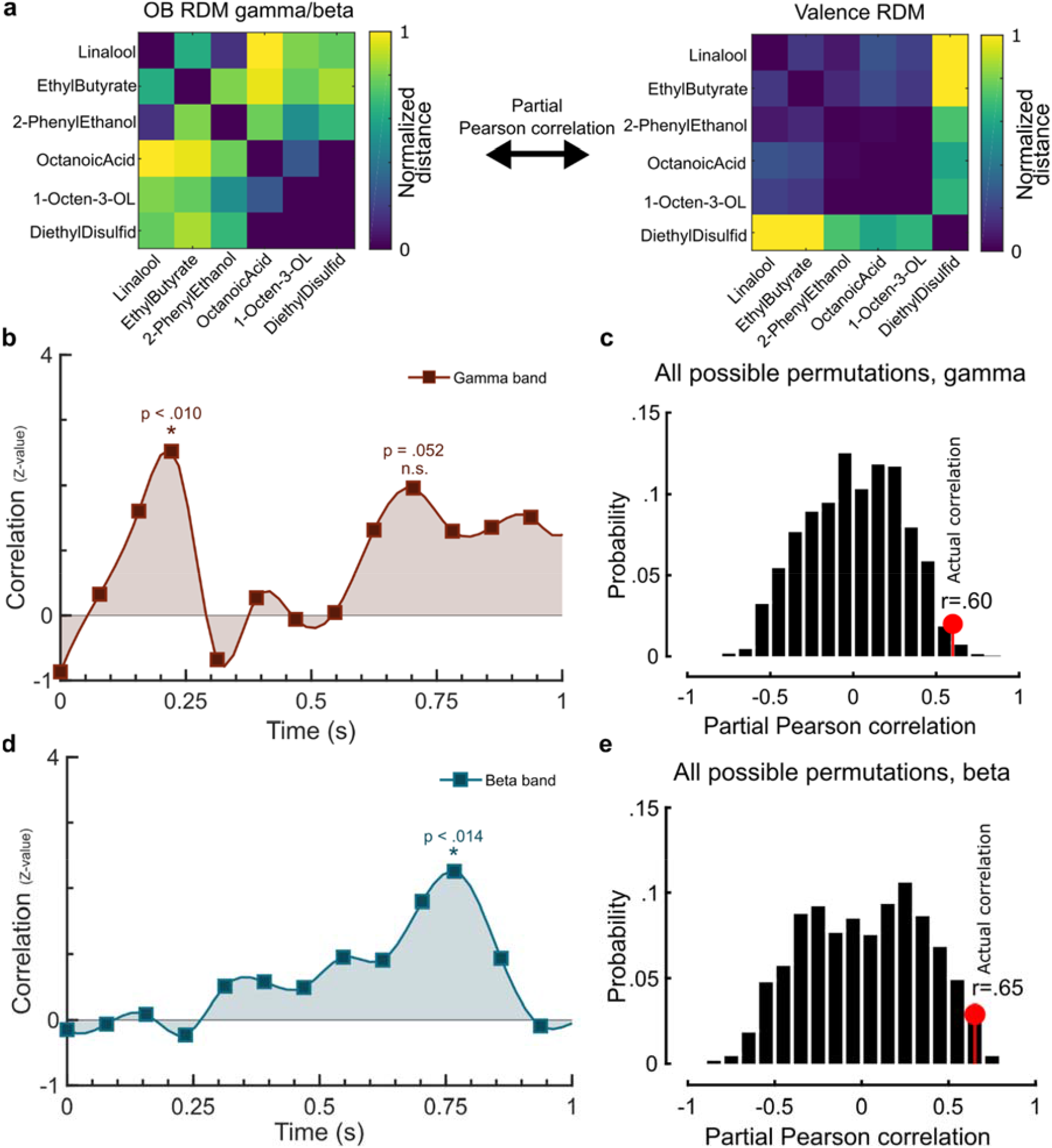
OB activity in the gamma and beta band relates to valence perception. **a**) Example of relationships between valence and OB activity in RSA **space b**) Partial Pearson correlation time course between activity of gamma and valence on the group level. All possible permuted partial Pearson correlation of gamma activity and valence indicated significant correlations at a time point ∼250-325ms after odor onset (*p* < .010). **c**) Distribution of all possible permutations for the significant instances and actual correlation indicated with red closed circles. **d**) Correlation time course of beta activity in the OB and odor valence RDMs indicating a significant relationship with valence perception around 800ms after odor onset. **e**) Distribution of all possible permuted partial Pearson correlation of beta activity and valence ratings for time points centered around 800ms (*p* <.014). Red closed circle shows the actual correlation within the permutation distribution.

Given the association between gamma activity and perceived valence, as well as the coupling between gamma amplitude and phase of beta in the t-PAC analysis, we subsequently assessed the potential relationship between beta band and perceived valence using RSA. The extracted time course of OB was band-passed to align with the results from the previously mentioned co-modulogram PAC (16-18 Hz). Similar to gamma, beta oscillation values were extracted and converted into complex signals using Hilbert transform to estimate instantaneous amplitude and phase values. This was next transformed to RDMs, and partial Pearson correlations were performed between beta RDMs and valence RDM, while controlling for perceived odor intensity, in a similar manner as described above.

We found that there was a significant association between beta activity and valence in a time-interval around ∼800ms after odor onset (**Figure 2d**). In other words, there was an association between perceived odor valence variance and beta activity variance around 800ms after odor onset in a sub-band around 16-18Hz. Similar to the gamma activity analysis, we tested the statistical significance of the correlations using all possible permutation tests for each time point within 1s after odor onset. Next, we compared each actual correlation with the distribution derived from the permutation to extract the exact p-value (*r* = .65, *p* < 0.014, *CI* = [.59, 1]). The distribution and exact *p*-value for the significant instance is shown in **Figure 2d**.

Behavioral studies in humans have demonstrated intensity-dependent regulation of the sniff response amplitude as early as 160ms after odor onset (Johnson et al. 2003) and there are demonstrated links between sniff magnitudes and both odor valence (Prescott et al. 2010) as well as odor intensity (Laing 1983). Similarly, sniff rhythms have been demonstrated to regulate OB gamma oscillation in anesthetized rodents (Manabe and Mori 2013). Therefore, to determine whether the discovered link between valence ratings and OB activity is potentially mediated by participants’ sniff patterns, unrelated to the odor presented, we assessed potential relationships between gamma and beta activity and relevant sniff parameters (sniff trace, i.e. amplitudes over time; max sniff amplitude; area under the curve) using separate Spearman rank correlations. However, our analysis demonstrated that there were no significant relationships between sniff trace and OB activity in either the gamma or beta bands (Supplementary **Figure S2**).

Next, we asked whether there were any commonalities in the neural representations of odor valence in the above identified gamma and beta bands. To this end, the group level distance matrices of gamma and beta (RDMs) at the identified time periods were scaled down using the first two eigenvectors (principal component, PC) into 2-dimensional (2D) representations. For the gamma band, valence seemed to be somewhat linearly organized along the first principle component (PC1) whereas there was no obvious valence-dependent organization along the second principal component axis (PC2) (**Figure 3a**). For the beta band, there was a reverse relationship as well as a linear linkage between valence organization between the PC1-PC2 dimensions. To investigate statistically which frequency band best explained most of the individual’s valence ratings, we first determined whether the organization of the odors within the 2D projections formed clusters. To this end, the RDMs were first converted to similarity matrices and communities were evaluated using a Newman algorithm (Newman and Girvan 2004). Odor valence ratings were hierarchically clustered from 1 to 6 clusters and we found the elbow of modularity index (*Q*) graph at 3, which indicate that a 3 cluster solution best explain valence ratings (**Figure 3b**). On the neural data, we subsequently derived a modularity index (*Q*) for each of the gamma and beta correlation peaks given the 3 clusters determined by hierarchical clustering of valence ratings and normalized their values to a corresponding null model from 5000 random re-wirings (Maslov and Sneppen 2002). We found that the beta band had a larger *Q*-value than the gamma band, meaning that the odors formed the most coherent pleasant and unpleasant clusters here. This indicates that more detailed information of participants’ subjective valence ratings can be obtained from the beta frequency band than the gamma band (**Figure 3c**). We then statistically assessed whether the obtained modularity indexes were significantly different from the null model using 5000 Monte-Carlo permutations tests. We found that the modularity index for beta was significantly larger than for gamma (*Z* = 2.95, *p* < .003, *CI* = [0.009, 0.018]).

**Figure 3.**
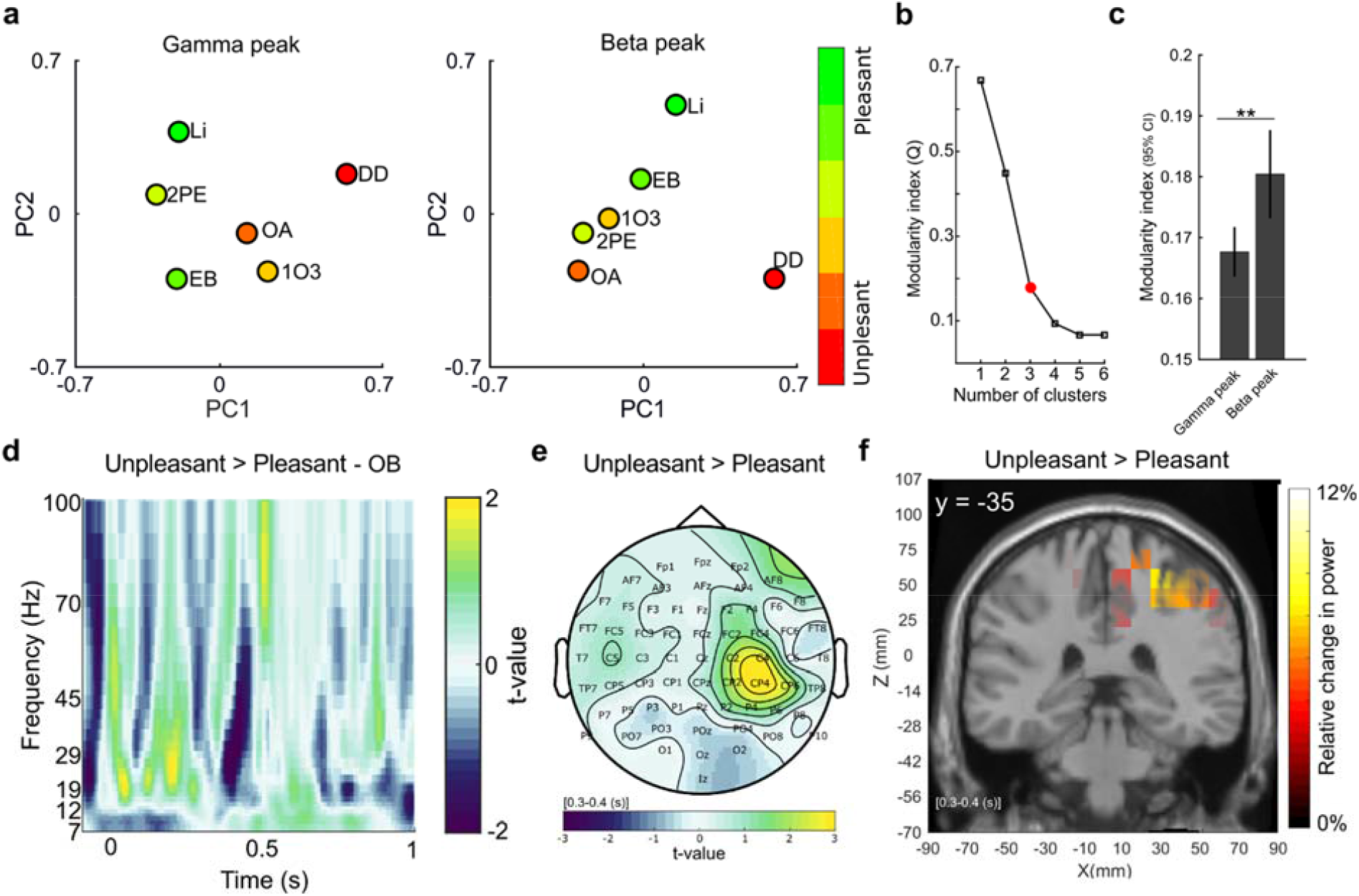
OB activity in the late-beta band is more similar to subsequent perceptual ratings than activity in the early gamma band. **a**) Odors placed within the 2-dimensional PC space, derived from peak values within detected significant peaks in Figures 1b &d, separated by frequency band. Observe the linear alignment between perceived odor valence and placement in PC space for the beta band. Odor names are written using abbreviations: Li (Linalool), 2PE (2-phenyel Ethanol), EB (Ethyl Butyrate), 1O3 (1-Octen-3-OL), and DD (Diethyl Disulfide). Colors code indicate average perceived valence where green colors denote positive valence and red/yellow colors denote negative valence. See Figure 1b for absolute valence ratings. **b**) Hierarchical clustering valence rating from 1 to 6 clusters. The elbow of the graph is shown with a red closed circle. **c**) Newman modularity index demonstrates that mean Q-values are larger for beta synchronization indicating that a more detailed odor valence readout can be inferred from this time point. Error bar shows 95% confidence interval for 5000 permutations and ** in panel C indicates p < .01. **d**) t-contrast map indicated more beta power during early, and less beta power during late time points, for the two most unpleasant odors compared to two most pleasant odors. **e**) Topographical map of mu rhythm illustrated higher values for unpleasant compared to pleasant odors over motor cortex during interval of 300-400 ms. **f**) Source of mu rhythm was localized to right motor cortex (x 27, y -35, z 60) using eLORETA during the time-interval displayed in **e**), i.e. 300-400 ms after odor onset.

These results suggest that the final odor valence perception can best be explained by processing in the beta band. Our analyses so far have, however, assessed odor valence by forcing ratings into a single continuous dimension or into three clusters and used these continuous parameters to assess organizational relationships between odor perception and neural activity. This approach means that we cannot assess whether either one of the contrasting valence dimensions (pleasant or unpleasant) contribute more to the OB processing. It has been argued that positive and negative valence is separated in a 2-dimensional space (Schiffman 1974) and a common approach in past studies has been to assess valence using a dichotomized design where groups of odors that differ in their rated valence (labeled as pleasant and unpleasant) are contrasted. To facilitate an assessment of whether there are differences in processing between pleasant and unpleasant odor in the OB, we compared OB processing of the two most pleasant odors against the two most unpleasant odors, eliminating the two neutral middle odors, all based on the individual’s own valence rating. When contrasting the two odor valence categories, we found that negative odors produced a greater synchronization response in the early portion of the beta band (around 50 to 200ms, *t* = 3.01, *p* < .004, *probability CI-range* = .004) whereas positive odors produced a greater synchronization response in the late beta band (around 690 to 780ms, *t* = 3.49, *p* < .002, *probability CI-range* = .003) determined by 5000 Monte Carlo permutation tests.

The separation in processing between the two valence extremes demonstrated that negative odors produced a more pronounced activity in the early beta band. We hypothesized that this early processing might be indicative of the initiation of an early avoidance response. This would fit behavioral data in humans that have shown that an odor associated with threat elicits a full body motor avoidance response (Arshamian et al. 2017). If this hypothesis is valid, we should observe odor valence-dependent modulation of preparatory motor responses over motor cortex in the time-interval of these OB responses. Specifically, we hypothesized that we would observe greater power in the mu rhythm over the motor cortex for negative odors. Desynchronization in mu rhythm has previously been demonstrated to be a measure of preparatory motor responses to salient stimuli (Kumar et al. 2013; Matsumoto et al. 2010), whereas inhibition of motor behavior yields synchronization in mu rythm (Howe and Sterman 1972; Pfurtscheller et al. 2006). To this end, the mu rhythm for two extremes was assessed on the whole scalp where we found greater power over motor cortex (electrode C2: *t* = 2.17, *p* < .014, *probability CI-range* = .003; electrode C4: *t* = 3.00, *p* < .003, *probability CI-range* = .001; electrode CP2: *t* = 2.01, *p* < .022, *probability CI-range* = .004; electrode CP4: *t* = 3.27, *p* < .001, *probability CI-range* = .001; electrode CP6: *t* = 2.23, *p* < .012, *probability CI-range* = .003). Next, we localized the source of the mu rhythm using eLORETA and found a cluster around the right motor cortex (x 27, y -35, z 60) where the dipole voltage density was 12% stronger for unpleasant compared to pleasant odors. Furthermore, for each trial, we extracted valence ratings and odor induced responses and subsequently assessed effect on mu rhythm for each trial using general linear model with Valence and Intensity as predictors. The group effect of Valence to predict mu power was finally estimated for each electrode using the student t-test. In-line with our hypothesis, odor valence was related to mu rhythm power over motor cortex in the time-period of interest (250-450ms after odor onset, the interval was selected slightly larger to increase frequency specificity), electrode CP2: *t*(17) = -2.20, *p* < .042, *CI* = [-1.46, -0.03] and electrode FC4: *t*(17) = -2.25, *p* < .037, *CI* = [-.96 -.03] (Supplementary **Figure S3**). In other words, the more negative an odor was perceived, the more mu power over the motor cortex was observed in the time period of the early OB processing.

### Unpleasant odors elicit a fast avoidance response

Our results so far have suggested that processing in the human OB is attuned to odor valence and that there is a link between processing in the OB and motor cortex activity in a valence-dependent manner. These results suggest a link between valence processing of negative odors in the OB and an early avoidance response. In other words, if results obtained in Experiment 1 are valid, when a negative odor stimulates the OB, a behavioral avoidance response should be initiated by the motor system in the time period shortly after the demonstrated mu activity, i.e. 400ms plus motor response time. To directly test this prediction, in Experiment 2, we sought to determine whether odor valence initiates an approach-avoidance response. Specifically, we wanted to determine whether this response is initiated based on processing in the early time period, where associations between OB processing and valence perception were found as well as a functional gating between the OB and motor cortex in the mu band. We operationalized approach-avoidance motor responses as posterior-anterior angular motion, derived from normalized responses from a force plate that measures participants’ whole-body micro-sway (**Figure 4a**). We hypothesized that a negative odor would elicit an avoidance response, manifested by the initiation of a backward motion in the early time period of interest. The body micro-sway was measured as posterior-anterior angular motion that was normalized to the height of individuals and band-pass filtered to produce posterior-anterior momentum (PAM). Two pleasant and two unpleasant odors, with averaged valence rating illustrated in **Figure 4b**, were presented using sniff-triggered olfactometry, identical to what is described for Experiment 1.

**Figure 4.**
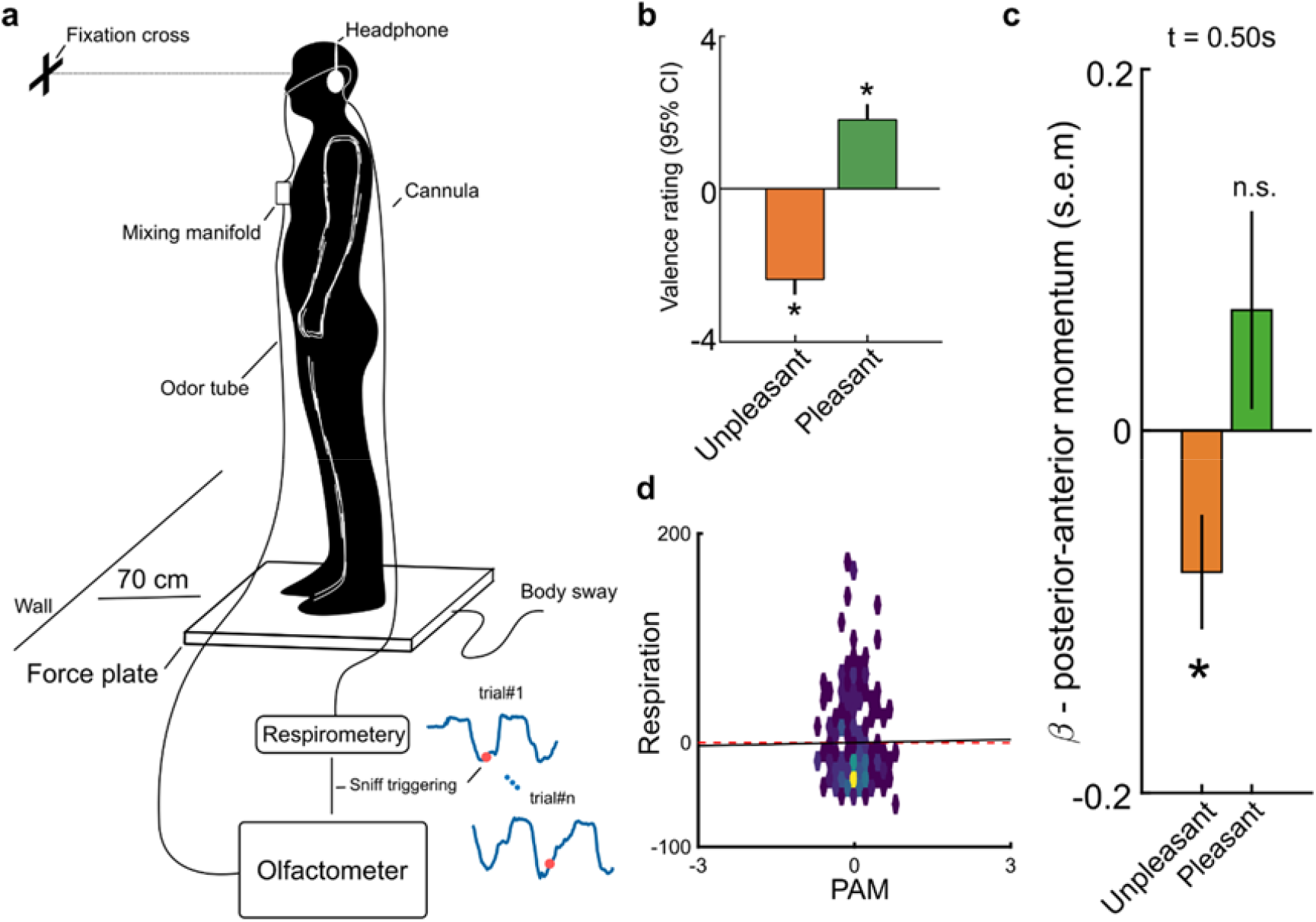
Unpleasant odors elicit a fast avoidance response. **a**) Experimental setup in pilot and Experiment 2. Participants stood centrally on the force plate, with their feet together, facing a wall with a fixation cross placed at their eye level. Continuous respiration was measured using a respirometer and the olfactometer was triggered close to the nadir of a respiration cycle to synchronize the trial onset with inhalation. **b**) Bars show averaged valence rating of unpleasant and pleasant odors during the experiment (error bars showing 95% confidence intervals). **c**) Valence-dependent modulation of PAM was replicated in the main experiment in line with our preregistrered hypothesis and indicating significant backward movement (i.e. beta values below zero) for unpleasant odors 500ms after odor onset. **d**) Non-significant correlation coefficient between PAM and respiration flow suggest that differences in breathing did not facilitate differences. Heatmap shows the joint distribution and dashed red line shows the correlation. Star in figures denote *p* < .05.

We first performed a pilot experiment (n=21) to allow us to determine time-point(s) of interest for analyses of PAM responses in a non-biased manner, to pre-register our hypothesis and analyses, and importantly, to establish known priors for subsequent Bayesian analyses. To this end, we assessed five time-points of interest, 0.25s (at the time of gamma processing), 0.5s (the hypothesized period of interest, gamma + response time, based on results in Experiment 1), 0.75s (at the time of the beta processing), and 1.0s (at the time of beta processing + response time) after odor onset across the pleasant and unpleasant odor conditions (**Figure S4a**). These time-points were selected to cover the full odor presentation with additional motor response time factored in. Within a LMM statistical model, with participant as intercept and conditions as random slope, we found a significant main effect for conditions only at the time point 0.5s after odor onset, *t*(61) = 2.13, *p* < .037, *CI* = [.01, 0.04], with no other significant effects at other time-points (**Figure S4b**). Subsequent Student’s *t*-tests against 0 (standing straight) demonstrated that the backward motion in response to negative odors was significant, *t*(61) = 2.06, *p* < .04, *CI* = [-0.28, -0.04], but without a potential forward motion in response to a positive odor, *t*(61) = 0.64, *p* > .74, *CI* = [-0.19, 0.39].

In the main experiment (Experiment 2; n=47), we focused our analyses on the time-point identified in the pilot experiment – all other aspects but the sample size were identical. We selected the sample size based on an estimated effect size of 0.3 (derived from the pilot experiment), required power 0.95, alpha error probability .05, and a correlation among measures of 0.4 – this yields a suggested sample size of 47 participants to enable a strong prediction. All hypothesis and analyses were pre-registered at https://aspredicted.org/fk9gw.pdf

We could replicate the result demonstrated in the pilot experiment with a significant difference between the two odor categories at time-point 0.5s after odor onset, *t*(174) = 3.24, *p* < .001, *CI* = [0.06, 0.23] (**Figure 4c**). Given the fact that for Experiment 2 we had a known prior from an independent dataset (result from the pilot experiment), we further explored this effect using Bayesian statistics. The Bayesian analyses supported results obtained with frequentist methods. We found that our analyses gave substantial support for a difference between the two parameters (Bf10 = 3.32; Supplementary **Figure S5**). However, these analyses assess potential differences between the two odor categories whereas results from the pilot experiment indicate that effects are mediated mainly by negative odors. When assessing each odor valence category separately against no movement using one-sided Student’s *t*-test with directionality based on the pre-registration, there was once again only a statistically significant effect for the negative odors to elicit a backward motion, *t*(174) = 2.47, *p* < .007, *CI* = [-∞, -0.016], and no significant effect for the positive odors, *t*(174) = 1.22, *p* > .11, *CI* = [-0.04, +∞].

We then assessed whether the difference between odor valence in PAM was mediated by a potential difference in respiratory flow, a parameter that previously has been demonstrated to be linked to odor valence (Frank et al. 2003). However, we found no correlation between the respiration flow and PAM at the time point of interest (0.5s), *rho* = - 0.02, *p* >.75, (**Figure 4d**). Moreover, to verify the lack of dependency, we assessed the null effect using Bayesian statistics with the known priors. These analyses also supported the conclusion that PAM and respiration were not interdependent (Supplementary **Figure S6**).

## DISCUSSION

We here demonstrate that neural activity in the human olfactory bulb (OB) is linked to perceived odor valence. Specifically, we found that the OB processes odor information sequentially within two time periods; first a brief period of initial gamma activity across the valence dimension, and a temporally privileged early beta activity for unpleasant odors, both indirectly linked to the initiation of a motor avoidance response. Second, there was a later period of beta processing that was linked to the linear formation of the final subjective valence percept of the presented odor. These results indicate that one of the initial and primary functions of the OB is to process early odor-based valence information; potentially to extract early odor-based warning signals.

The observation that odor valence was processed in the human OB mainly during two time-points, one early and one late stage, is in-line with the two-stage model of odor processing in the OB suggested by Frederic and colleagues (2016). They specifically demonstrated that the OB executes a first fast processing, relying on gamma oscillations, allowing the individual to make fast discriminations, and a second slower processing, relying on beta processing, utilizing information from centrifugal inputs to support more deliberate decisions. Our data, which is in line with earlier human and animal work, suggests that the OB processes valence of unpleasant and pleasant odors at different time points. In this context, our link between early gamma and beta (for unpleasant odors) oscillations in the OB suggests that this might be a preparatory, non-deliberate mechanism for fast avoidance of potentially dangerous odors or avoidance of their sources. This hypothesis is further supported by previous findings that odor-induced gamma oscillations within the OB are largely a local phenomenon (Martin and Ravel 2014; Gray and Skinner 1988) and sometimes dependent on the individual’s behavioral state or past negative experience with the odor (Kay 2003). Moreover, because the OB is located very early in the odor processing stream with projections from the olfactory receptor neurons and monosynaptic connections to cerebral areas associated with processing of information related to threat/saliency (Mainland and Sobel 2006), the OB is ideally localized in the processing pipeline to process avoidance related information given the need for a fast response. Indeed, associating an odor with an aversive outcome alters a range of parameters in how the OB processes the associated odor. For example, it increases neural responses (Kay and Laurent 1999; Fletcher 2012) as well as axonal density into associated glomeruli, which in turn is associated with an increase in associated glomeruli size (Jones et al. 2008). Based on our past findings, it is therefore likely that one of the initial and important aspects of the OB is to extract and process odor information that is associated with a potential threat.

Results from the modularity index, derived from the principal component analyses, indicated that the strongest link between the two time periods and final odor valence rating was the later beta activity occurring around 700-800ms after odor onset. This is also in line with the two-stage model (2016) of odor processing in the OB with a separation of processing between bottom-up and top-down dependent processing. Beta oscillations are often considered more of a ‘top-down’-dependent signal in opposition to gamma that is considered as more of a ‘bottom-up’-dependent signal (Gnaedinger et al. 2019; Richter et al. 2017; Kay 2014); although, it should be made clear that beta has been demonstrated to be initiated in the OB during odor sampling (Gourévitch et al. 2010). Past studies in animal models have indeed linked gamma band processing within the OB to intra-bulb processing (Kay 2014; Martin and Ravel 2014) and beta processing has been demonstrated to sometimes be dependent on centrifugal feedback from piriform cortex (Bressler 1984; Martin et al. 2006) and to be modulated by context or past odor associations (Frederick et al. 2016). Our data cannot, however, conclusively determine whether the OB activity in the later time period originates from centrifugal information from higher order areas. Nonetheless, we believe that the most parsimonious explanation is that the late beta is representing valence-dependent signals that project back, directly or indirectly, from other cerebral areas, such as the orbitofrontal, amygdala, and piriform cortex. These projections would then help shape the final interpretation of the odor by adding information of past experiences (Li et al. 2008) as well as information about the odor object per se (Wilson and Sullivan 2011; Porada et al. 2019; Olofsson et al. 2014) — two factors that are known to strongly influence the final odor valence percept (Yeshurun and Sobel 2010; Djordjevic et al. 2008).

In our data, the early gamma and beta power in the OB seems specifically attuned to the processing and perception of negative odor valence with no clear demonstration that the OB processes odors associated with a positive outcome in early time points. This finding is in-line with recent findings in rodents indicating that odors with a positive valence are processed initially within an area downstream from the OB, the olfactory tubercle (Gadziola et al. 2015; Gadziola and Wesson 2016), an area that does not have centrifugal direct connections with the OB (In ‘t Zandt et al. 2019). Whether this regional-specific valence separation in processing also occurs in humans is not known. However, a recent study demonstrated that intracranial electric stimulation of the OFC, an area previously linked to odor valence processing in humans, could only produce pleasant odor experiences (Bérard et al. 2020). The separation demonstrated in rodents does, however, support the 2-dimensional valence hypothesis that odor valence perception is not represented by a unidimensional spectrum, ranging from unpleasant to pleasant. Instead, a 2-dimensional space where positive and negative valence is separated with neutral valence as the start point has been suggested (Schiffman 1974). This has also been supported by data on the semantic distribution of odor descriptors that are commonly divided into a positive and a negative category (Schiffman 1974; Schiffman et al. 1977). In contrast to the demonstrated increase in beta frequency for odors with more negative valence as compared to odors with more positive valence, we did not find an association between valence and beta activity when assessing links in RSA space using a linear approach in early time points. A possible explanation for this discrepancy might be that our t-PAC finding suggests that the detailed odor valence information for beta is reflected in phase and not amplitude in early time periods of OB processing of the odor. Hence, it is possible that the lack of linear association between neural activity in beta band and odor valence in RSA space during early time points is due our finding that only phase, but not amplitude, based on PAC coupling, is linked to gamma, thereby suggesting that the coupling of amplitude is critical for finding linear association in RSA space. Similarly, our present method may not be able to capture that the OB also processes positive valence during early time points due to the range of our odors used where only some mean individual ratings reached above 70 on the 0-100 scale (see **Figure 1b**). It should further be noted that the EBG method requires that participants are in a nutrition deprived state to maximize signal from the OB, which might have an impact on the obtained results (Iravani et al. 2020; Royet and Pager 1981; Pager et al. 1972). However, this later aspect of the method can also mean that the potential perceived reward of the odor would increase and therefore also maximizes the likelihood of finding effects for positive odors. Future studies where odors are either individually selected based on their reward properties or conditioned with positive outcome are needed to conclusively determine whether the human OB prioritizes processing negative odor valence.

Results in Experiment 1 suggested the existence of an approach/avoidance motor response occurring around 500 ms after odor onset in a valence-dependent manner. In two separate experiments (pilot and Experiment 2) and using a design with preregistered hypotheses and analyses, we demonstrate here that odors with a negative valence triggered an avoidance response that was manifested as leaning away from the odor source. We have previously demonstrated a similar fast avoidance response in human participants to the odor of blood (Arshamian et al. 2017), an odor that is treated as an approach or avoidance trigger across species. Our current findings extend these results and suggest that odor avoidance in humans might extend beyond biologically important and potentially inherent signals and be a general phenomenon that is linked to the valence of the odor per se. Negative odors induced larger mu desynchronization over and within the motor cortex which is a response that previously has been demonstrated to be a measure of preparatory motor responses to salient stimuli (Kumar et al. 2013; Matsumoto et al. 2010). Although speculative, this mu desynchronization appeared in a time period that, when response time is factored in, corresponds to the motor avoidance response to negative odors demonstrated in Experiment 2. Moreover, this motor cortex response to the negative odors appeared around 150ms after the valence-related increase in gamma and early beta activity within the OB, thereby allowing the signal time needed to transmit the information between the OB and motor cortex. For neutral and pleasant odors, a similar motor movement would probably be related to later processing. In line with this notion, rodents preforming a Go, No-Go task with positive reinforcement exhibit a simultaneous activity in the piriform cortex, primary motor cortex, and the mRN just before executing a motor response for the Go trials. Importantly, while these responses are seen in low and high gamma, as well as beta activity, they are only present during the second and third (final sniff) (Hermer-Vazquez et al. 2007). For the first sniff (equivalent to the data in our study), there was, however, no response related to the motor cortex for the rewarded GO trials. This indicates that for neutral and pleasant odors, as compared to unpleasant, a top-down regulation from olfactory cortex and motor cortex directly to the OB is more evident (Hermer-Vazquez et al. 2007). Our results suggest that the olfactory and motor systems are more closely linked also in humans than has previously been appreciated and that this, especially for unpleasant odors, may be initiated at the OB-level.

It should be noted that while our results demonstrate that the human OB processes subjective odor valance, it does not suggest that the OB is the first processing stage of valence. Multiple studies in humans and non-human animals alike show that an odorant’s valence partially depends on its physicochemical properties (Khan et al. 2007; Secundo et al. 2014; Poncelet et al. 2010; Bensafi et al. 2007; Mandairon et al. 2009; Joussain et al. 2011; Keller et al. 2017). Physicochemical properties have been shown to predict olfactory receptor neuron activation and based on this, it has been suggested that odor valence is, at least partially, coded at the level of the olfactory epithelium (Joussain et al. 2011). Because the 6 odors in Experiment 1 were selected from the DREAM challenge (Keller et al. 2017) to span the physicochemical valence space, our OB activations may to some extent reflect information from physicochemical properties originating from receptor neurons projecting upstream to the OB (Sezille et al. 2015). It is still an open question if a peripheral valence code originating from physicochemical properties reaches the OB, and if the OB in turn refines the signal or keeps it unchanged. It should, however, be noted that all results linking odor valence to OB processing is based on the individual’s own valence rating and not an a priori defined valence rank among the included odors. Specifically, we show that perceived pleasantness is represented uniformly across participants in the bulb at specific time periods and frequencies in a manner that is represented by subjective perception and not pre-defined odor classification. This suggests that valence is processed and not merely manifested by odor identity. Valence ratings is, to a not trivial degree, dependent on personal experiences. Indeed, when we assess the relationship between individual’s RDM of odor valence ratings and that of the full group, the mean similarity is 61.6%, which means that about 38.2% of the total variance in these valence ratings (in this case, valence ranking) is explained by individual differences. Our choice of methods therefore reduces, but do not eliminate, the potential impact physicochemical properties might have on our results in favor of subjective valence perception.

Even though we can demonstrate links between the OB and motor cortex in a relevant time-period, it should be noted that our whole-body avoidance results are only indirectly linked to the EEG data. For it to be directly linked, it would require assessing EEG source signal from individuals that are freely moving around which is not possible with current method for measuring OB responses because the active-electrodes strongly amplifies motion artifacts. To the best of our knowledge, no method currently exists that would allow measures from the OB while participants moved their full body. Moreover, it is important to highlight that our data only covers the first sniff of an unannounced odor. As demonstrated in rodents, OB processing is continuously updated with each continuous sniff with marked shifts in both neural and behavioral responses (Gupta et al. 2015; Patterson et al. 2013; Hermer-Vazquez et al. 2007).

In summary, our results suggest that the human OB processes odor valence. We propose that the two stages of processing of valence in the OB is due to a reciprocal process where the initial fast gamma and beta response address negative odors, valence information that may be projected from the olfactory receptor neurons (Khan et al. 2007) or learnt from past aversive experiences already coded in the OB (Kay and Laurent 1999). In contrast, the later beta response seems more related to the final valence rating of the odor as well as potential preparation of the OBs initial gamma and subsequent beta responses for the second sniff of the same odor. At this time stage the OB should be influenced by information related to past experiences with the identity of the odor. Importantly, negative odors seem to have a privileged temporal access in the human olfactory bulb. This suggests that one of the initial functions of the OB is to process and extract early odor-based warning signals to aid the individual’s approach-avoidance decisions.

## METHOD

### Experiment 1 – Valence decoding from oscillations within the olfactory bulb

#### Participants

In Experiment 1, 19 individuals (mean age 28.88 ± 4.52, 7 women) who reported being healthy, non-smokers, and with no history of head trauma or neurological disorders, participated in three separate recording sessions (all identical). Prior to inclusion, a working sense of smell was confirmed in all participants using a 5 item, 4 alternative, cued odor identification test (Kobal et al. 2000). All participants cleared the cut-off for inclusion of at least 3 correct answers. Given the scarcity of functional anosmia in the participants’ age-range, the probability that we erroneously included individuals with anosmia in the experiment is less than 0.05%. The study was approved by the local ethical review board (EPN: 2016/1692-31/4) and all participants signed informed consent prior to participation.

#### Chemicals and odor delivery

In Experiment 1, the six odorants that were used came from the DREAM Olfaction Prediction Challenge(Keller et al. 2017). In the DREAM project, participants rated odor pleasantness of 476 structurally and perceptually diverse molecules. We averaged ratings across all participants in the DREAM project and ordered odors from the most pleasant to the most unpleasant. Based on this ranking, we selected six that evenly spanned the entire valence dimension. Next, we conducted a pilot study on Swedish participants and verified that the odorants elicited similar ratings in a Swedish context and diluted them down to iso-intense concentration. Odors used were: 0.14% Linalool (Sigma Aldrich, CAS 78-70-6), 0.25% Ethyl Butyrate (Sigma Aldrich, CAS 105-54-4), 0.1% 2-Phenyl-Ethanol (Sigma Aldrich, CAS 60-12-8), 0.2% 1-Oceten-3-OL (Sigma Aldrich, CAS 3391-86-4), 1% Octanoic Acid (Sigma Aldrich, CAS 124-07-2), and 0.25% Deithyl Disulfide (Sigma Aldrich, CAS 110-81-6), all diluted in neat diethyl phthalate (99.5% pure, Sigma Aldrich, CAS 84-66-2) and dilution values are given in volume/volume from neat concentration. Lack of trigeminal sensations in their presented concentration was established using odor laterality tests (Hummel 2000; Wise et al. 2012).

Odors were delivered birhinally using a computer-controlled olfactometer and each odor was presented 20 times in each session to participants (i.e., 60 times in total for each odor across the three sessions). The olfactometer has an onset-time of 200ms, measured from computer trigger to odor delivered in the nose, and a sharp rise-time to facilitate an odor presentation with high temporal precision (Lundström et al. 2010). A total flow rate of 3 liters/minute (l/min), inserted into a constant 0.3 l/min flow to prevent tactile sensation of odor onset was used. To further avoid that participants could predict odor onset, yet assure a clear percept, odor onset was (unbeknown to the participants) triggered by their own sniff cycle. When the assigned inter-trial-interval (10s) had occurred, an odor was triggered at the nadir of the inhalation phase in the sniff cycle following that interval, thus assuring odor presentation at inhalation. A relatively long inter-trial-interval was used to lower the risk of odor habitation. Participants sniff-cycle was measured by thermopod (Experiment 1) and respirometer (Experiment 2) sampling at a rate of 400 Hz (Powerlab 16/35, ADInstruments, Colorado) and respiration traces for triggering of odor were analyzed on-line by the LabChart recording software (ADInstrument, Colorado). Data were subsequently down-sampled off-line to 40 Hz and processed in MATLAB 2018a for further analyses.

E-prime 2 (Psychology Software Tools, Pennsylvania) was used to record event timing, triggering, and sniff data logging. Jittered pre-stimulus interval of 600∼2000ms was placed before each trial to minimize anticipation of odor onset. Participants were tested in a sound attenuated booth, purpose amended for odor testing, with good air circulation to minimize potential redundant disturbances. Moreover, during the experiment, low volume white noise was presented to participants via headphones to mask potential airflow-related auditory cues from the olfactometer that might possibly cue odor onset.

#### Procedure

To allow us to collect a large data set for each individual, each participant participated in 3 sessions on separate days with at least one day and at the most a month apart. Each session consisted of three 15 min long blocks with 5 min break between each to limit odor adaptation/habituation, totaling about 1h per session. Participants were presented with the 6 different odors in a random order and after each odor presentation, they rated how pleasant and intense they perceived the presented odor to be. Ratings were done by placing a marker on a labeled visual analogue scale presented on the screen ranging from 0 (very unpleasant / very weak) to 100 (very pleasant / very strong).

#### Electroencephalography, Electrobulbogram, neuronavigation measurement

Sixty-four EEG scalp electrodes were placed according to international 10/20 standard and an additional 4 EBG electrodes on the forehead (Iravani et al. 2020). Signals were sampled at 512 Hz using active electrodes (ActiveTwo, Bio-Semi, Amsterdam, The Netherland). Electrode offsets were manually checked prior to experiment onset and electrodes were adjusted until meeting the a priori established criteria (<40mV). Next, the position of all the electrodes in stereotactic space were determined using an optical neuro-navigation system (Brain-Sight, Rogue Research, Montreal, Canada); for more detail please see (Iravani et al. 2020).

#### EBG/EEG Data Analysis Preprocessing

Data were epoched from 500 ms pre-stimulus to 1500 ms post stimulus, re-referenced to average of activity at all electrodes, linear phase band-pass filtered at 1-100 Hz (Butterworths 4^th^ order), and power-line interference filtered using discrete Fourier transform (DFT) filtering at electrical frequency (50 Hz) to remove power line noise. Trials with large muscle and eye blink artifacts were identified with an automatic algorithm. The artifact detection protocol comprised of band-pass filtering using Butterworth filter (4^th^ order), Hilbert transform to extract amplitude values, and z-scoring. Trials with z-values above 4 were marked and removed from further analysis.

#### Olfactory bulb time course extraction

To extract OB’s time-course, digitized electrode positions were first used to co-register participants head to a default MNI brain using a six parameters affine transformation. Second, a head model was constructed based on a multi-shell spherical head model. Spherical volume conductors were considered for scalp, skull, gray matter, and white matter with the conductivity of 0.43, 0.01, 0.33, and 0.14 (Iravani et al. 2020). The covariance matrix of electrodes during the 1s odor presentation were regularized by 10% prior to be fed into extra low-resolution electromagnetic tomography (eLORETA) algorithm to estimate the time-course of the dipole placed in (x ±6, y 30, z -32) on trial level, which corresponds to OB location (Pascual-Marqui et al. 2011; Iravani et al. 2020). The maximum projection of the dipoles’ time course over three principal axes were computed to serve as OB activity. eLORETA analysis was carried out in the open source Fieldtrip toolbox 2018 within MATLAB R2019b (Oostenveld et al. 2011).

#### Time-frequency analysis of olfactory bulb signal

After extracting the OB’s time-course, we assessed the difference in power evolution of the two most unpleasant and pleasant odors. The time-frequency map for broadband frequencies [1∼100 Hz], with step of 1 Hz and interval [-0.1∼1 s] with step of 0.005s, was estimated using multi-tapered sliding window from discrete prolate spheroidal sequences (DPSS). The window length was adjusted to cover at least 3 cycles for each frequency, ranging [.3∼3s]. Next, the time-frequency map of each trail was assessed and converted to decibels. Finally, to create the contrast map, the most unpleasant and pleasant category, each consisting of two odors, were determined on the individual level based on participants’ valence ratings and contrasted against each other.

#### Mu rhythm and source localization

Similar to time-frequency analysis of OB signal, the mu rhythm power for all scalp electrodes were estimated using multi tapered sliding DPSS widow for mu frequency range [10 ∼13 Hz] with step of 0.5 Hz and time interval of [.3 ∼ .4 s] with step of .005 s. Likewise to OB time-frequency analysis, the window lengths were chosen to cover at least 3 cycles of mu rhythm. Next, the mu power for the two most unpleasant and pleasant odors were estimated, baseline corrected and converted to decibels. Finally, a topographical map was created and non-parametric statistics were performed to create contrast and find channels that were significantly different in mu power. Source localization were performed similar to the olfactory bulb source localization, and after co-registration of electrodes to default MNI brain, a spherical head model with 4 tissue type was created. The cross spectral density matrix of electrodes during the .3 ∼.4s after the odor onset were regularized by 10% and fed into eLORETA to localize the source of mu rhythm.

#### Time resolved phase-amplitude coupling (t-PAC)

t-PAC of the extracted OB time-course were analyzed between gamma and beta bands with window length 250ms and 50% overlapping. The gamma band [30∼100Hz], discretized to 20 frequency bins, and the instantons amplitude were extracted using Hilbert transform at each frequency bin. Similarly using Hilbert transformation, the instantaneous phase of slower oscillations (i.e. beta [12∼30Hz]) were computed and t-PAC for each time bin was calculated as the power ratio of the composite signal of instantaneous amplitude of faster and phase of slower to the faster oscillation during the window interval on the individual level (Samiee and Baillet 2017). Moreover, we quantified the co-modulogram level between the outcome of t-PAC with a slower band to isolate a range of slower frequencies that are coupled to identified gamma. t-PAC analysis were carried out in BrainStrom toolbox within MATLAB R2019b (Tadel et al. 2011).

#### Representational similarity analysis (RSA)

RSA is a powerful tool to connect different levels of experimentations (e.g. behavioral and brain data) beyond linear correlation. Here, we used RSA to assess the relationship between the odor valence rating and neuronal population activity of OB in two prominent odor related frequencies, gamma and beta. To limit our statistical tests and minimize the potential false positive error, we only include the frequencies of gamma and beta bands in the RSA analysis that were found to be coupled in the t-PAC analysis. Hence, the OB time-course was band-pass filtered using a Butterworth 5^th^ order to the (by the t-PAC) identified frequencies within the gamma (53-65 Hz) and beta (16-18 Hz) bands following a Hilbert transform to extract the instantaneous amplitude and phase. In RSA, instead of assessing the first-order isomorphism between the stimulus and brain representation, similarity in relationship within the stimuli and brain representation are assessed using representational dissimilarity matrix (RDM). To construct the RDM, OB Euclidean distances were used for averaged left/right OB activity and valence rating on the individual level. A temporal neighborhood with a radius of 7 samples (considering our sampling rate of 512 Hz, this equals to ∼14ms) was used, including both amplitude and phase, for fast gamma oscillations and 28 samples (equals to ∼55ms) for slower beta oscillations. The difference in neighborhood radius selection is due to the difference in temporal resolution and ensures at least one cycle of either faster or slower oscillation is included in the feature space. The RDMs for whole 1s of odor stimuli were compared between the behavioral and OB response in a searchlight framework on the individual level. In line with a multidimensional scaling method (Abdi et al. 2005), the so-called DISTATIS method, we constructed a consensus RDM to represent the group level. To determine the potential relationship between neural and perceptual RDMs, values above diagonal line of the matrices was assessed using all possible permuted (i.e. shuffling the labels of odors; given 6 odors, the total possible combinations is 720) partial Pearson correlation to avoid inflated correlation due to symmetry of RDMs (**Figure 1a**). The group level RDMs were subsequently scaled down using eigenvector decomposition into two main axes. The distance matrices were converted to similarity matrices by inversing the distance matrix after added by 1, and modularity indices (Q) were computed using the Newman method (Newman and Girvan 2004) given three clusters. The three clusters were identified using a hierarchical clustering of valence rating, varying the number of clusters from 1 to 6 clusters and estimating the knee of the modularity curve where the knee of the curve was estimated as the furthest point from the linear approximation. RSA analysis and community detection were performed in the open source CoSMoSMVPA toolbox (Oosterhof et al. 2016) and MATLAB Network Toolbox https://github.com/ivanbrugere/matlab-networks-toolbox.

#### Statistical analysis

We assessed the statistical difference in power evolution between the two most unpleasant and pleasant odors, as well as scalp mu rhythm, using a non-parametric statistic. The time-frequency maps of unpleasant and pleasant odors of OB signal and scalp electrodes were computed using multi-tapered sliding windows and compared using 5000-permuatation Monte Carlo tests to find significance time/frequency bins or channels. To statically test the relationship between the brain data and valence rating in RSA, RMD matrices at each time point were shuffled through all possible combinations. In each iteration, partial Pearson correlation between neural and perceptual valence was computed with intensity as nuisance covariates. To extract the exact p-value from the permutation test, we computed the number of times the actual partial Pearson correlation was bigger than shuffle data out of total permutations (720). Similarly, for t-PAC analysis, non-parametric Monte Carlo 5000 permutation tests were performed for the OB coupling value at each time-frequency bin against baseline (250ms pre-stimulus) and exact p-value were extracted. Subsequent t-map was smoothed, while preserving the shape, for illustration purpose.

The distance matrices at the instances of significant correlation with valence for both gamma and beta were scaled down to first and second principle components (i.e., PC1 and PC2) and Newman modularity was calculated to assess whether the tested odors clusters comply with valence rating. To statistically test the Newman index Q, the modularity of similarity matrices was compared with the null model for each correlation peak. The null model was generated by 5000 times rewiring of the adjacency (similarity) matrix while persevering weight, degree and strength distribution using Brain Connectivity Toolbox within MATLAB R2019b (Rubinov and Sporns 2011; Rubinov and Sporns 2010). Later the actual modularity index was compared with the null distribution. In the post-hoc analysis to assess effects on pre-motor responses, we extracted data in the time interval 300-400ms after odor onset. We tested if the preparatory response for motor action we observed in Experiment 2 could be found in this experiment, even though here participants were sitting on the chair and instructed to sit as still as possible. The power of mu synchronization/desynchronization was predicted using a generalized liner model having valence, and intensity as predictor for each electrode on the scalp and yielding in beta maps on the individual level. Later on the beta maps were statically tested on the group level via the student t-test.

### Experiment 2 – Odor valence-dependent approach/avoidance responses

#### Participants

Given that links between perceived odor valence and approach/avoidance motor responses had not been previously assessed, we initially performed a structured pilot experiment to explore the time period of interest of the motor response, the result of which was later used as a priori defined temporal regions of interest in Experiment 2. In the pilot experiment, a total of 21 individuals (age = 28.71 ± 5.84, 11 women) participated. In the subsequent Experiment 2, a total of 47 individuals (age = 25.94 ± 4.2, 29 women) participated. Inclusion criteria (including passing the anosmia screening test) were the same as described above for Experiments 1. The studies were approved by the local ethical review board (EPN: 2016/1692-31/4) and all participants signed informed consent prior to their participation.

#### Odors and delivery method

In both the pilot and the main experiments, odors were piloted and presented as described in Experiment 1. In the pilot experiment, 4 odors were used; namely 10% Strawberry (IFF Inc.), 50% Carvone (CAS 6485-40-1), 50% Fish odor (IFF Inc.), and .0014% Ethanethiol (CAS 75-08-1). To limit odor dependency, Strawberry was substituted with vanillin, 2.55 grams in 12 milliliters (CAS 121-33-5), and Ethanethiol with 0.25% Diethyl Disulfide (Sigma Aldrich, CAS 110-81-6) in the main experiment. However, the strawberry odor was after the experiment deemed to be contaminated and was subsequently removed from all analyses in the pilot experiment; thus, in the pilot experiment, only responses to 3 odors were analyzed. All odors were diluted in neat diethyl phthalate (99.5% pure, Sigma Aldrich, CAS 84-66-2) and concentration values were volume to volume except for vanillin that was available in crystal form. Odor selection and concentration was based on previous pilot experiments where a range of odors were rated for perceived valence and after sub-selection, concentration-adjusted to achieve intensity matching.

Identical to Experiment 1, odors were (unbeknown to the participants) triggered by their sniff cycle, measured by a respirometer (**Figure 4a**) sampling at the rate of 1000 Hz (Powerlab 16/35, ADInstruments, Colorado), subsequently down-sampled off-line to 40 Hz and processed in MATLAB 2018a for further analyses. A relatively long inter-trial-interval of 10s was used to reduce the risk of odor habitation.

#### Body sway measurement

Participant’s body micro-sway was assessed with a force plate (AccSway^Plus^, AMTI Massachusetts) assessing 8 axes of motion. The force plate was initially allowed to warm up for a few minutes after which it was zeroed and a 25 seconds period of unloaded baseline was initially recorded for calibration purposes.

#### Procedure

Participants stood in the center of the force plate with their feet together, facing a wall where a fixation cross was placed at eye-height about 70 cm away from their face. The height of the fixation cross was adjusted for each individual according to participants’ height. Their arms were positioned alongside the body and they were instructed to avoid performing redundant movement (**Figure 4a**). Odors, with 20 repetitions, were presented in blocks with 4 odors, in each using a pseudorandom order where the individual randomization order was adjusted so that no odor was presented consecutively over the 80 presentations. After each odor presentation block, an auditory cue was presented via headphones to cue participants to provide a verbal rating of the perceived odor intensity of the odors presented within the preceding block. This task was performed to force participants to focus on the odor but not specifically their perceived valence. A total of 4 blocks, each 5 minutes long, was presented. To prevent potential compliance effects, participants were told that the aim was to assess the relationship between sniff and odor intensity and first after the experiment they were informed about the true aim. At the very end, participants were asked to rate odor valence of each odor (**Figure 4b**). This was done at the end to prevent participants from discovering the true aim of the experiment and potentially bias their response in the desired direction.

#### Statistical analyses

Posterior-anterior angular momentum, extracted from the body sway data recorded by the force plate, was assessed according to a calibration matrix provided by the force plate vendor. In addition, as suggested by the vendor (AMTI, Watertown, USA), the offset correction is implemented by removing the average of 25 seconds unloaded recording (i.e. without participant standing on the force plate) prior to each session from the experimental force plate data. Extracted posterior-anterior angular momentum was subsequently normalized to the individuals’ height to estimate the linear posterior-anterior momentum (PAM). These PAM parameters were then epoched to [0, 1.25s], linear and quadratic trend were removed and band-pass filtered using linear phase FIR filter ([0.45 ∼ 2 Hz], n=93, hamming window, - 53dB attenuation of stopband) to remove normally occurring micro-sway, produced by the motor system to sustain balance.

In the pilot study, one individual was removed from analyses after rating the unpleasant odors as very pleasant (>3 standard values from the mean), thus leaving the total as n=21. Later, event related responses of the PAM were calculated for five time points during the odor interval with steps of .25 second to identify the time point of interest. Linear mixed effect model (LMM) with by participant intercept and random slope of odor were fitted for the three time points. Having determined the time point of interest, we repeated the experiment in a completely new data set with bigger sample size and fitted LMM with exact similar design. Moreover, as a control analysis we examined if respiration correlates PAM by means of Pearson correlation at the time point of interest.

The analysis was repeated in a Bayesian framework as supplementary analysis. For the pilot experiment we considered a normal distribution with mean of 0 and standard deviation of .5 and half Cauchy with standard deviation 5 for random effect (Supplementary **Figure S5a**). For control analysis, correlation was modeled with multivariate normal distribution with means of μ1 and μ2 corresponding to mean of respiration and PAM as well as a covariance matrix Σ containing variances and correlation value for respiration and PAM. Prior distribution for means was normal distribution with mean of 0 and standard deviation of 20, prior for variances was uniform distribution with range of [0, 50] and for correlation a uniform distribution with range of [-1 1].

## ACKNOWLEDGEMENT

We thank Kimberly Battista (battistaillustration.com) for the making of EBG insert in Fig. 1A. Funding provided by the National Institute on Deafness and Other Communication Disorders (R21DC016735) as well as the Knut and Alice Wallenberg Foundation (KAW 2018.0152), awarded to JNL. AA is supported by a grant from the Swedish Research Council (2018-01603)

