## Supplementary Figures for "The human olfactory bulb process odor valence representation and initiate motor avoidance behavior"

---

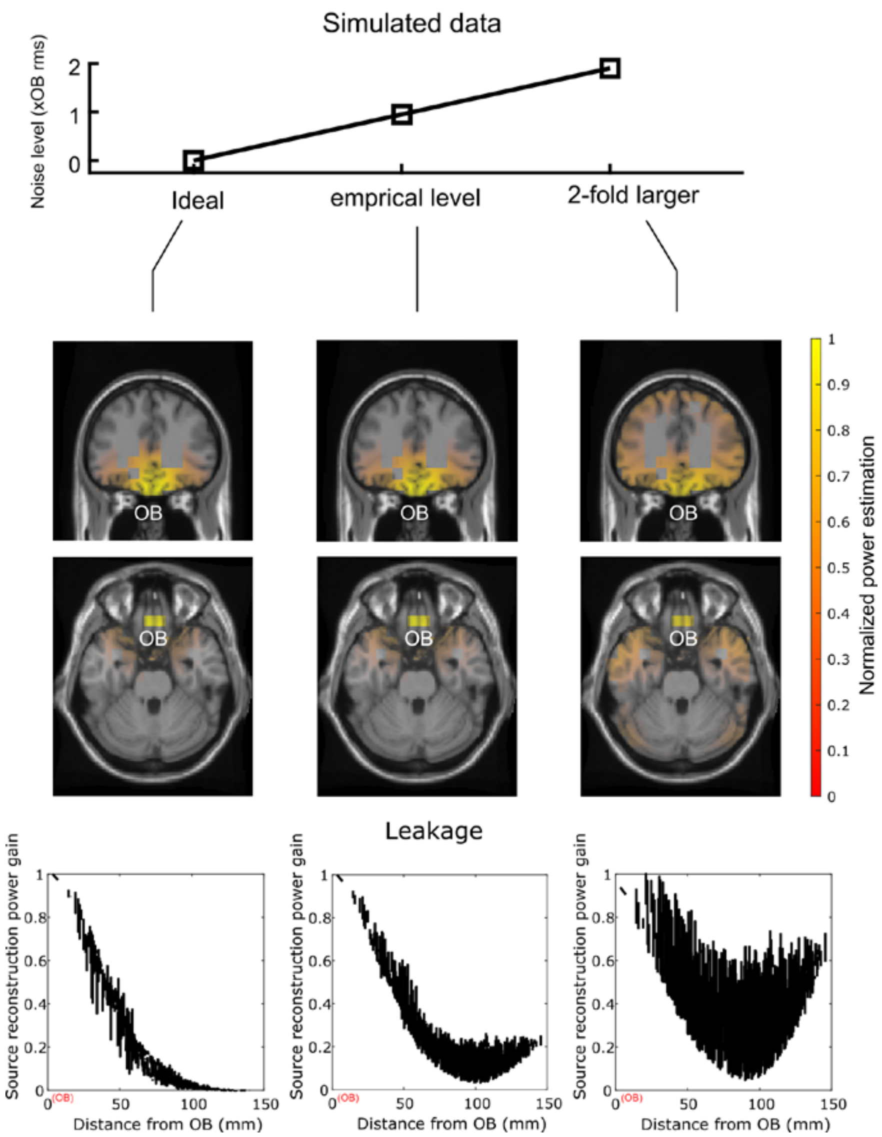


***Figure S1. Simulating eLORETA spatial dispersion for OB source.*** *Three models were tested, the ideal, the empirical, and a model with 2-fold larger noise level; all shown in upper panel. Middle panel shows eLORETA reconstruction spatial gain for the three levels of noise. Lower panel shows the eLORETA gain (proportional to leakage) as function of physical distance from OB.*

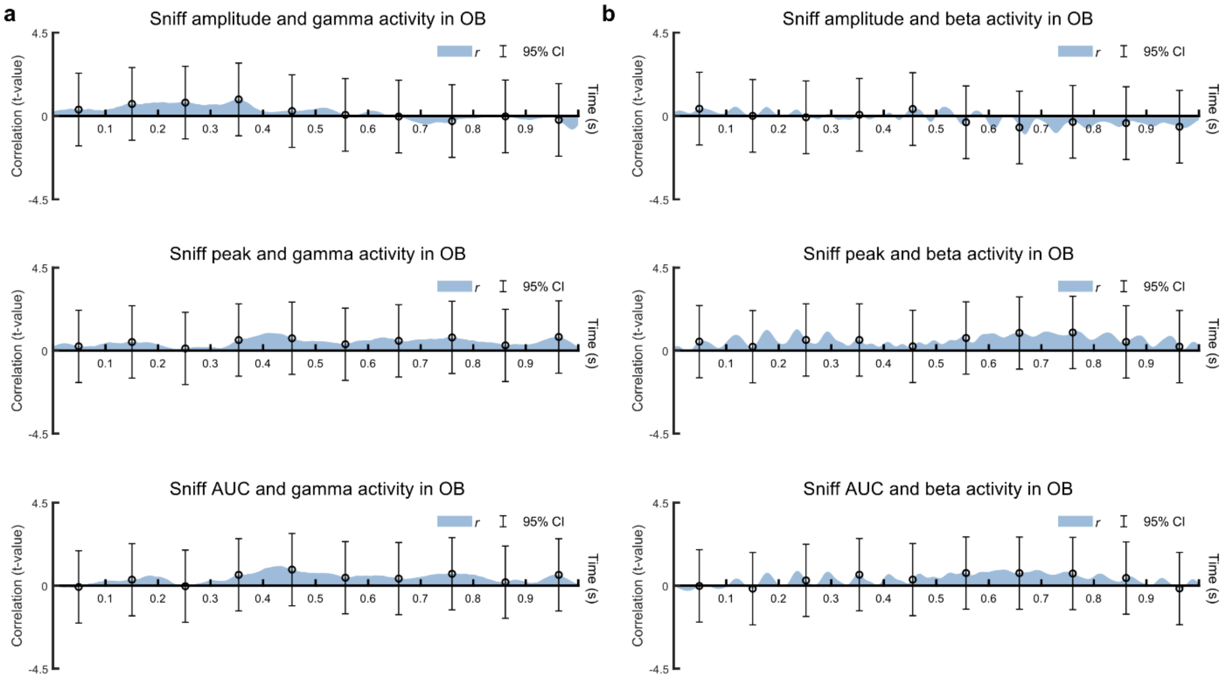


***Figure S2. Gamma, beta activity and sniff association a****) Person correlation between gamma activity and sniff parameters including amplitude, peak and area under the curve (AUC) of the sniff and gamma activity in OB showed no dependency between sniff and gamma activity.* ***b****) No dependency has been found between beta activity and sniff parameter such as amplitude, peak and area under the curve (AUC) using Pearson correlation.*

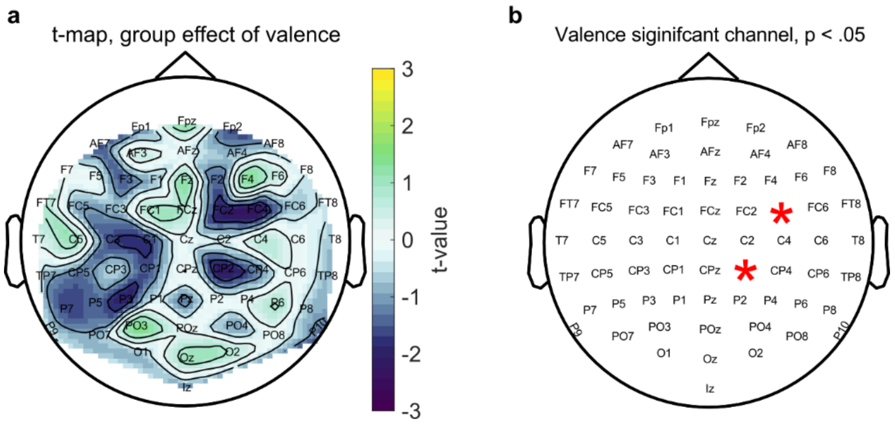


***Figure S3. Relationship between mu desynchronization and rated valence. a****) t-map shows the effect of valence on the scalp mu power. Color bar indicate t-values and each marking on the scalp is an electrode position according to the international 10/20 convention.* ***b****) Red stars over electrode names indicate significant electrodes.*

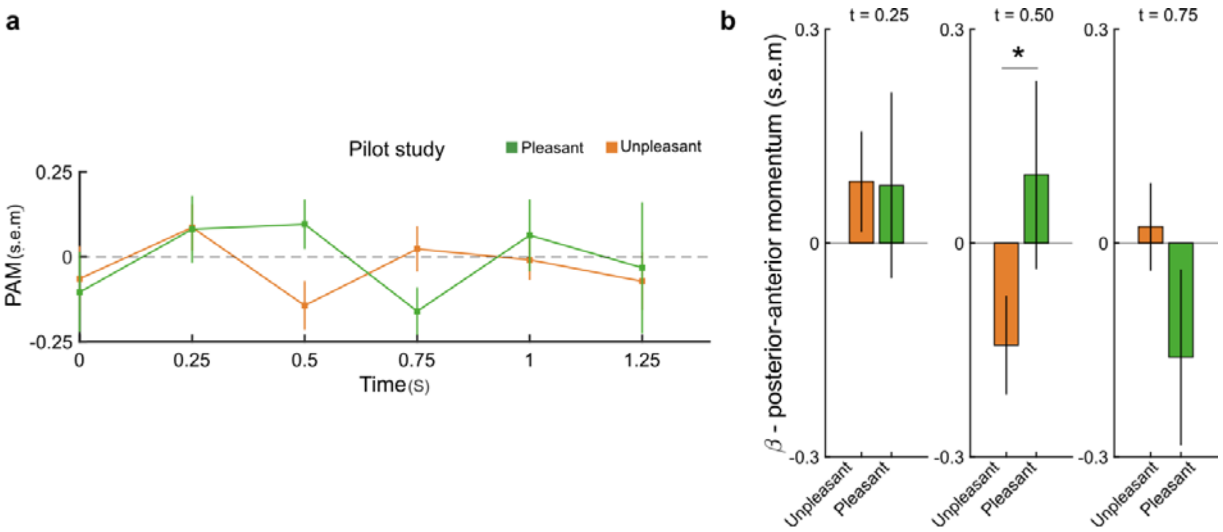


***Figure S4. Pilot experiment to test different the time points for approach-avoidance response. a**) posterior-anterior momentum (PAM) as function valence across time. Squares shows the average of PAM for each time point and errorbars show standard error. **b)** Linear mixed effect model in the pilot study suggests that there is significant difference between PAM as function of valence at the time point of 0.5s. Bar graphs show the estimated beta value and error bars show standard errors.*

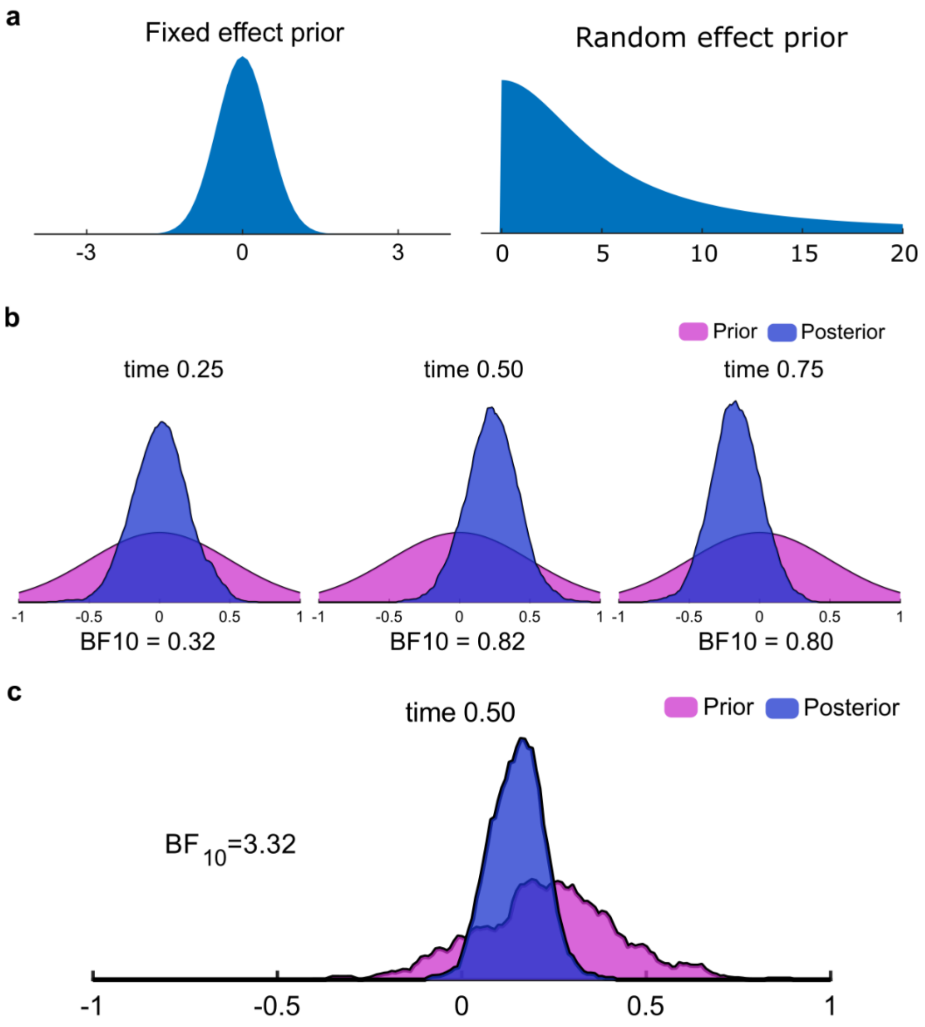


***Figure S5. Bayesian statistics for Experiment 2. a****) Prior distributions for the pilot experiment. Normal distribution with mean of 0 and standard deviation of 0.5 is considered for fix effect and half Cauchy with standard deviation of 5 for random effect.* ***b****) Prior and posterior distributions for three time points for the pilot experiment suggest a potential effect at time point 0.5s determined by highest* $BF_{10}$. ***c****) Prior and posterior for Experiment 2 at the time point 0.5s. The posterior (result) of the pilot experiment was used as the prior for analyses of Experiment. Our results (BF10>3.3) indicate that there is an effet of valence on posterior-anterior momentum (PAM) at the selected time-point.*

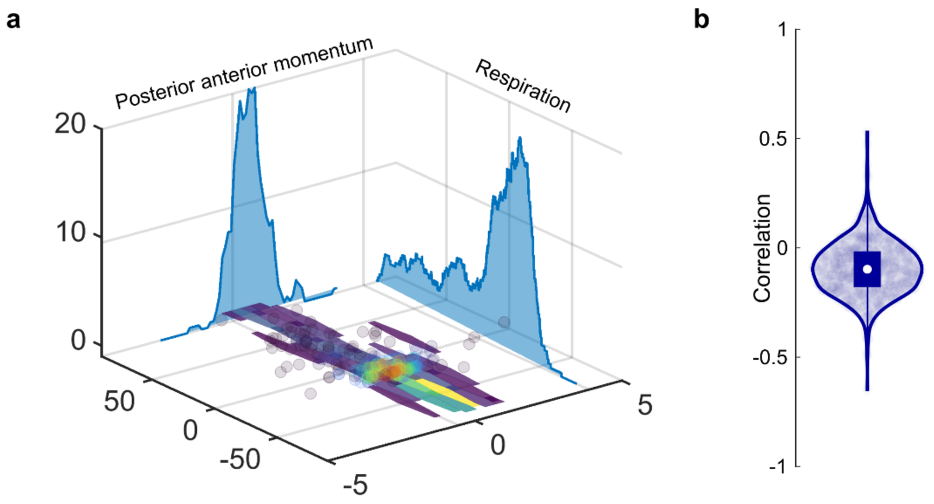


***Figure S6. Bayesian correlation between PAM and respiration in Experiment 2. a****) Two marginal distribution for posterior anterior momentum (PAM) and respiration is shown on the side planes (xz and yz). Scatter plot shows the underlying data and plotted on xy plane as well as heatmap that show the density of data points.* ***b****) Violin plot of the posterior distribution of correlation between PAM and respriation. White dot shows the median of distribution, box shows the interquartile range, whiskers show maximum and minimum values, and blue circles shows individual values.*
